# Early antiviral CD4 and CD8 T cell responses and antibodies are associated with upper respiratory tract clearance of SARS-CoV-2

**DOI:** 10.1101/2023.10.25.564014

**Authors:** Sydney I. Ramirez, Paul G. Lopez, Farhoud Faraji, Urvi M. Parikh, Amy Heaps, Justin Ritz, Carlee Moser, Joseph J. Eron, David A. Wohl, Judith S. Currier, Eric S. Daar, Alex L. Greninger, Paul Klekotka, Alba Grifoni, Daniela Weiskopf, Alessandro Sette, Bjoern Peters, Michael D. Hughes, Kara W. Chew, Davey M. Smith, Shane Crotty, the ACTIV-2/A5401 Study Team

## Abstract

T cells are involved in protective immunity against numerous viral infections. Data regarding functional roles of human T cells in SARS-CoV-2 (SARS2) viral clearance in primary COVID-19 are limited. To address this knowledge gap, samples were assessed for associations between SARS2 upper respiratory tract viral RNA levels and early virus-specific adaptive immune responses for 95 unvaccinated clinical trial participants with acute primary COVID-19 aged 18-86 years old, approximately half of whom were considered high risk for progression to severe COVID-19. Functionality and magnitude of acute SARS2-specific CD4 and CD8 T cell responses were evaluated, in addition to antibody responses. Most individuals with acute COVID-19 developed SARS2-specific T cell responses within 6 days of COVID-19 symptom onset. Early CD4 T cell and CD8 T cell responses were polyfunctional, and both strongly associated with reduced upper respiratory tract SARS2 viral RNA, independent of neutralizing antibody titers. Overall, these findings provide evidence for protective roles for circulating SARS2–specific CD4 and CD8 T cells during acute COVID-19.

## Introduction

Since the emergence of severe acute respiratory syndrome coronavirus 2 (SARS2) as a novel human pathogen, much has been learned about protective immunity to SARS2, both in the context of infection and COVID-19 vaccines. Serologic correlates of protection have been established for vaccines (1–8). In the context of prophylaxis, virus neutralization by monoclonal antibodies (mAb) has been demonstrated as one mechanism of protective immunity (9–12). In contrast, in the context of infection the relative importance of immune system compartments may differ, due to the substantial differences in kinetics of primary versus memory immune responses and lack of pre-existing antibodies. Studies of primary adaptive immunity to acute SARS2 infection offer a key opportunity to evaluate early humoral and cellular immune responses and their individual contributions to protection.

The global population rapidly developed widespread cellular and humoral immunity to SARS2 as a result of both vaccination and infection. However, key gaps in our understanding of human primary immune responses to SARS2 remain. Cellular immunity may be required for viral control and clearance during acute infection (13–17). While acute and memory T cell responses clearly occur to SARS2 infection (18–24) and COVID-19 vaccines (25–32), evidence of functional protective roles of T cells have been limited in humans (16, 33), particularly by assessing non-hospitalized COVID-19 cases (34–36). Due to the combined difficulties of identifying early acute COVID-19 cases, recruiting those individuals, and the technical challenges of measuring virus-specific T cell responses and viral loads concomitantly, most acute T cell studies have been limited to small cohorts, including hospitalized cases, sometimes relatively late in disease, and often without viral load measurements (22, 37–40). In a controlled SARS-CoV-2 human challenge study of young adults with a mean age of 22 years old, early circulating SARS2-specific CD8 T cells were associated with SARS2 viral clearance in that cohort of 18 subjects (36). There is evidence for cellular immunity contributing to SARS2 protection in non-human primate models (16, 17). Breakthrough infections represent a different context for studying protective immunity in humans, in which there is evidence for contributions of memory T cells (41, 42). Given expanded interest in next-generation COVID-19 and pan-sarbecovirus vaccines with T cell-specific components (43, 44), or vaccines that are entirely T cell-based (30, 45, 46), future vaccine designs and clinical trial designs would benefit from better fundamental understanding of T cell protective immunity to COVID-19 in humans (30, 34).

## Results

Longitudinal data were collected for 95 individuals with primary SARS2 infection in 2020, prior to the availability of COVID-19 vaccines, with sampling including nasopharyngeal (NP) swabs for SARS2 RNA, serum, and peripheral blood mononuclear cells (PBMCs), all collected in the context of a randomized, controlled clinical trial (ACTIV-2/A5401, NCT04518410). Herein, serum antibodies, SARS2-specific CD4 T cell, and SARS2-specific CD8 T cell response measurements are reported, which is the largest and most comprehensive acute viral and antigen-specific immune response data set of its kind.

All participants were enrolled within 7 days from positive SARS2 testing and 10 days from COVID-19 symptom onset (21, 47). Individuals studied herein were among those randomized to receive either 700 mg of the mAb bamlanivimab or placebo (saline) intravenously (**Fig. 1A**) on study day 0 (median of 6 days post-symptom onset; **Fig. 1B**). The 46 bamlanivimab treatment and 49 placebo subjects were similar with respect to baseline characteristics such as age, biological sex at birth, risk for progression to severe COVID-19, and time post-symptom onset at randomization (**Fig. 1A and S1A**, and ref (21, 47)). Peripheral blood was collected prior to mAb or placebo administration on study day 0. NP swabs were collected from all 95 participants by trained study staff prior to treatment on study day 0, with SARS2 viral RNA was detectable in the NP swabs of nearly all individuals (89%) (**Fig. 1C**), and no significant difference in SARS2 NP RNA between the bamlanivimab (treatment) and placebo groups (**Fig. S1B**). SARS2 NP RNA at study day 0 was not associated with participant age (r = 0.16, p = 0.12, **Fig. S1C**). SARS2 NP RNA levels declined over time (**Fig. 1D**), as expected (48–50), and as previously reported for the full ACTIV-2/A5401 cohort (47).

**Figure 1.**
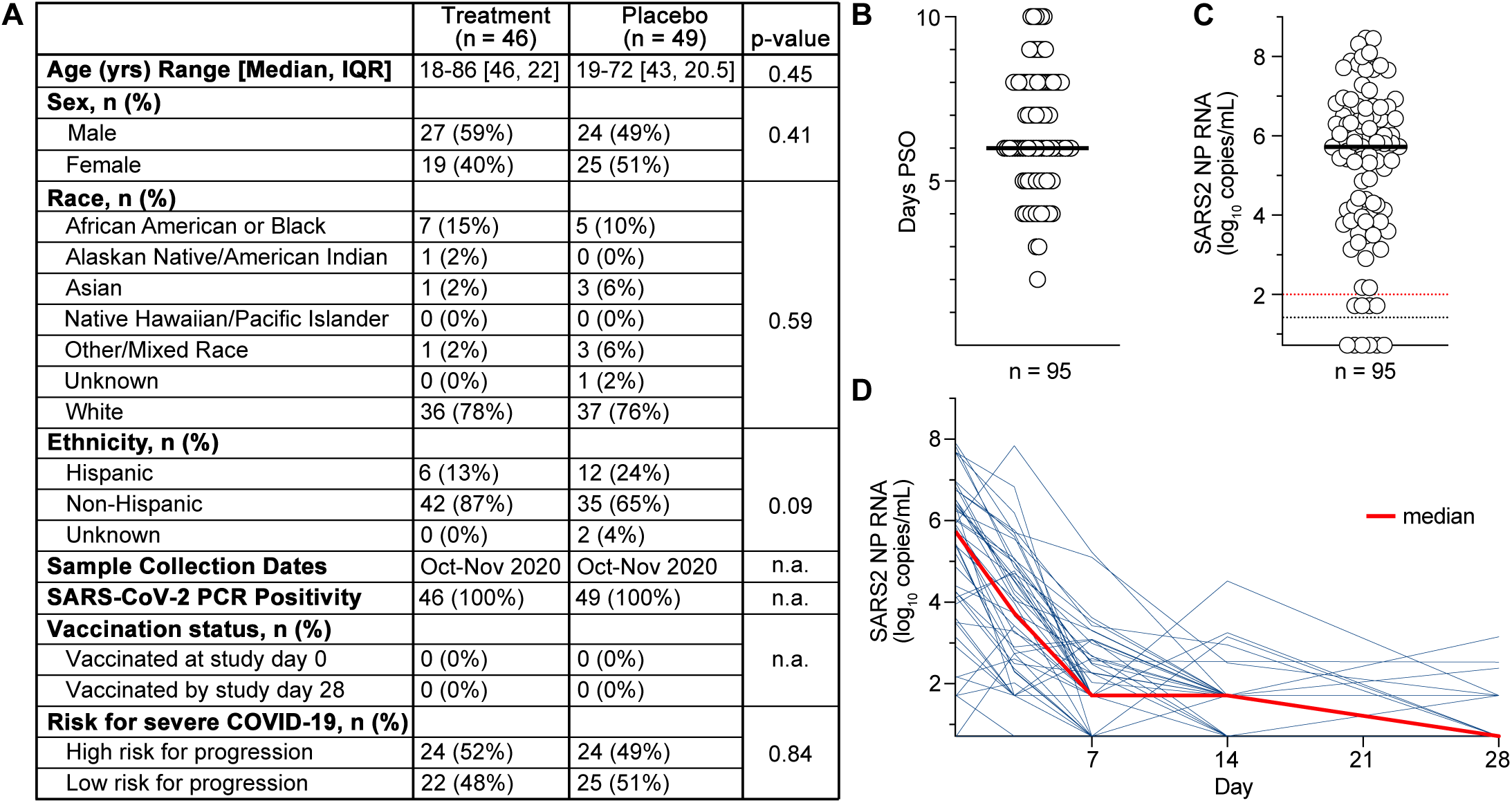
Study cohort characteristics and SARS2 NP RNA levels during acute COVID-19 and longitudinally. **A.** Demographics and other pertinent characteristics of the matched bamlanivimab (Treatment) and placebo group participants included in this study. Total n = 95 subjects. See methods for additional details. **B.** Median days post-symptom onset (PSO) from start of COVID-19 symptoms to study entry (study day 0) for all participants. **C.** SARS-CoV-2 NP RNA by quantitative reverse transcription polymerase chain reaction (RT-qPCR) for all participants prior to treatment on study day 0. Dotted black line limit of detection (LOD). Dotted red line limit of quantification (LOQ). Values >LOQ were considered positive. Bar = median. **D.** Longitudinal SARS-CoV-2 NP RNA data for the placebo group (n = 49) for study days 0, 7, 14, and 28. Median for each time point in red.

Both NP swab and peripheral blood samples were collected prior to treatment on study day 0. Acute adaptive immune responses were measured for all participants prior to treatment. Additionally, longitudinal responses were measured in the 49 placebo group participants post-treatment on study days 7 and 28. We previously reported day 28 antibody, memory CD4 T cell, and memory CD8 T cell outcomes (21). Early (study day 0) SARS2-specific CD4 T cell response magnitude and functionality were evaluated by multiple techniques and using multiple SARS2 antigens. Activation induced marker (AIM), and hybrid AIM plus intracellular cytokine staining (AIM+ICS) T cell assays were employed (**Fig. 2A-B**, **Fig. S2A**). SARS2 Spike (S)-and non-Spike-specific CD4 T cells were measured by AIM assays following stimulation with S and non-S (CD4-RE (ref 28)) peptide megapools (MP). Two sets of AIM phenotyping surface protein marker pairs were utilized to assign AIM assay positivity (expression of surface OX40^+^ and 41BB^+^, **Fig. 2A**, and surface OX40^+^ and CD40L^+^, **Fig. S2B**). SARS2-specific CD4 T cell frequencies were comparable by AIM based on OX40^+^41BB^+^ or OX40^+^CD40L^+^ surface marker expression (**Fig. 2A and S2B**). Seventy-seven percent of individuals were positive for S-specific CD4 T cells, and 79% of individuals were positive for non-S CD4 T cells (**Fig. 2A**). Combined responses to SARS2 S-and non-S epitopes were also quantified to evaluate total antiviral T cell responses per subject (**Fig. 2A and S2B. Fig. S2C-G**). Overall, substantial CD4 T cell responses were observed.

**Figure 2.**
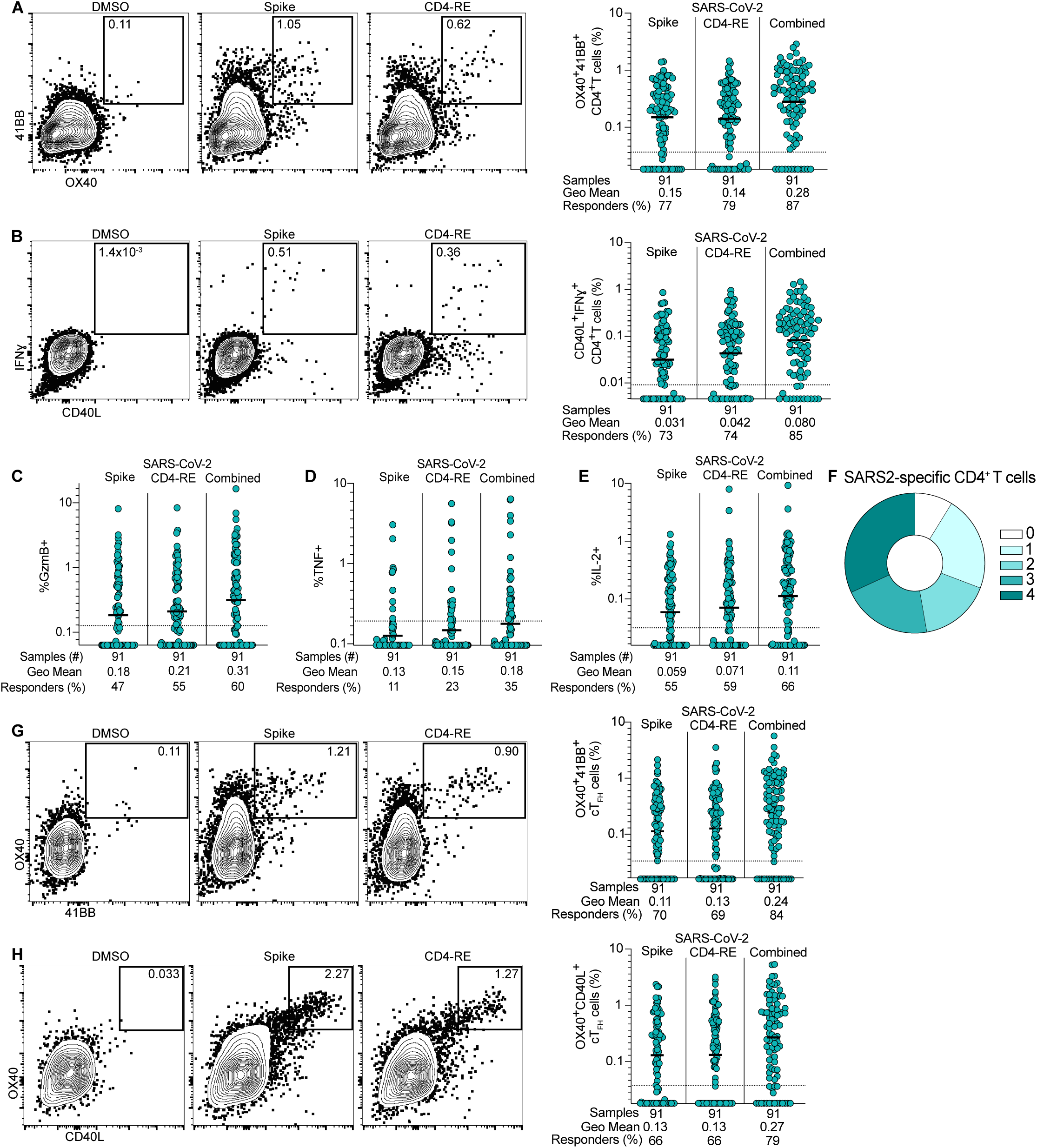
Antigen-specific CD4 T cell responses to primary SARS2 infection and acute COVID-19. **A-B.** Representative flow cytometry plots and frequency of SARS2-specific CD4 T cells to DMSO (negative control), Spike and CD4-RE MP stimulation conditions (Combined = sum of Spike + CD4-RE responses; see Methods for additional details) by (**A**) AIM using surface OX40 and 41BB co-expression, and (**B**) IFNγ ICS among surface CD40L^+^ cells. **C-E.** Study day 0 SARS2-specific CD4 T cell intracellular cytokine production (**C**) GzmB, (**D**) TNF, or (**E**) IL-2 among surface CD40L^+^ cells. **F.** Parts of a whole donut plot summary of intracellular cytokine (IFNγ, GzmB, TNF, IL-2) production by SARS2-specific CD4 T cells expressing 0 to 4 cytokines. **G-H.** As in **A** and **Fig. S2B** but for SARS2-specific circulating TFH (cTFH) cells. Bars = geometric mean. Dotted lines = LOQ. Flow cytometry gates display frequency (%).

SARS2-specific CD4 T cells were phenotyped based on cytokine production by ICS following stimulation with the S and non-S MPs, including interferon gamma (IFNγ), granzyme B (GzmB), tumor necrosis factor (TNF), and interleukin-2 (IL-2) production by CD40L^+^ CD4 T cells (**Fig. 2B-F and S2H-J**). IFNγ was the most expressed cytokine at study day 0 (**Fig. 2B**) and is known to be important in control of multiple viral infections in mouse models. Between 85-87% of individuals had SARS2–specific CD4 T cell responses by AIM or IFNγ ICS (combined S plus non-S responses; **Fig. 2A-B**). SARS2-specific CD4 T cells were polyfunctional, with 69% producing at least 2 cytokines, and 53% producing 3 or more cytokines (**Fig. 2B-F**). SARS2-specific circulating T follicular helper cells (cT_FH_, CXCR5^+^) were measured. Similar response rates were seen for SARS2-specific cT_FH_ as for total SARS2-specific CD4 T cells. SARS2-specific circulating T_FH_ cells were observed in 79-84% of participants’ study day 0 PBMC (**Fig. 2G-H**). Overall, SARS2-specific CD4 T cells with different functionalities and differentiation states were present.

In parallel, study day 0 SARS2 S-and non-S-specific CD8 T cell responses were evaluated by AIM (surface CD69^+^41BB^+^) and ICS following stimulation with SARS2 S and non-S (CD8-RE (ref 28)) MPs (**Fig. S3A, D-F**). Acute SARS2–specific CD8 T cell responses were observed at study day 0 in 63% of individuals by AIM (**Fig. S3B**). By ICS, 62% of individuals were positive for an acute SARS2–specific CD8 T cell response (surface CD69^+^ and intracellular IFNγ^+^, **Fig. 3A**). As was observed for SARS2-specific CD4 T cells, IFNγ was the cytokine most commonly produced by SARS2-specific CD8 T cells (**Fig. 3A**). CD8 ICS was used as the primary quantification of SARS2-specific CD8 T cells in downstream analyses given the similar detection rates of antigen-specific CD8 T cells but lower background for the CD69^+^IFNγ^+^ CD8 T cell assay (**Fig. 3A** LOQ) compared to CD8 AIM (**Fig. S3B** LOQ) (see Methods for additional details). As expected, CD8 T cell responses did not vary on study day 0 when assessed by clinical trial group assignment (**Fig. S3C**). Most CD69^+^ IFNγ-producing SARS2-specific CD8 T cells also produced GzmB (73%, **Fig. 3B and S3D**). Similar to SARS2-specific CD4 T cells, SARS2-specific CD8 T cells were predominantly polyfunctional, with 69% producing at least 2 cytokines and 58% producing 3 or more cytokines (**Fig. 3A-E**). Overall, polyfunctional virus-specific CD4 and CD8 T cell responses were detected in a substantial proportion of individuals during acute primary SARS2 infection.

**Figure 3.**
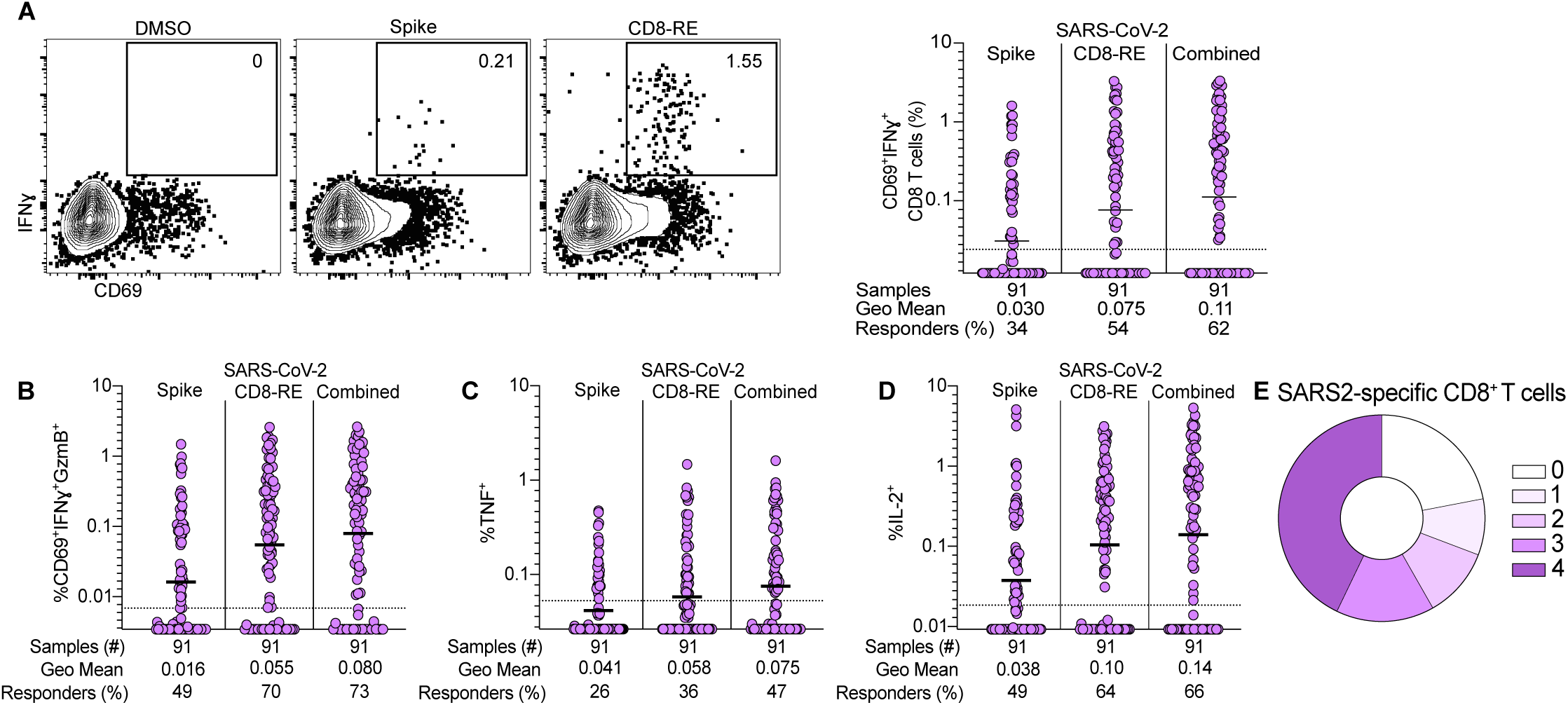
Antigen-specific CD8 T cell responses to primary SARS2 infection and acute COVID-19. **A.** Representative flow cytometry plots and frequency of SARS2-specific CD8 T cells to DMSO (negative control), Spike and CD8-RE MP stimulation conditions (Combined = sum of Spike + CD8-RE responses; see Methods for additional details) by IFNγ^+^ ICS among CD69^+^ cells. **B-D.** (**B**) IFNγ and GzmB, (**C**) TNF, (**D**) IL-2 production. **E.** Parts of a whole donut plot summary of intracellular cytokine (IFNγ, GzmB, TNF, IL-2) production by SARS2-specific CD8 T cells expressing 0 to 4 cytokines. Bars = geometric mean. Dotted lines = LOQ. Flow cytometry gates display frequency (%).

We examined the kinetics of T cell and antibody responses in the placebo group in more detail. Longitudinal assessment of SARS2-specific CD4 and CD8 T cell frequencies on study days 0 (**Fig. 2-3**), 7 (**Fig. 4A-C and S4A**), and 28 (**Fig. 4A-C**, **S4A**, and ref.(21)) showed that AIM^+^ CD4 T cell frequencies and the proportion of positive responders increased over time (**Fig. 4A and S4A**). SARS2-specific IFNγ^+^ CD4 and CD8 T cell frequencies remained stable from study days 0 to 28 (**Fig. 4B-C**), suggesting that antigen-specific IFNγ-producing T cells are formed early during primary infection and stably maintained at least through early convalescence. SARS2 serologic assessments, included nAb titers to ancestral SARS2 by lentiviral pseudovirus neutralization assays (PSV NT50) and receptor binding domain (RBD) binding IgG titers measured on study day 0 (**Fig. 4D-E** and ref 21 for RBD IgG). Most individuals were found to have formed nAb to SARS2 within 6 days post-COVID symptom onset. On study day 0, 62% of participants had positive nAb titers (**Fig. 4D**, **Fig. 4SB**). As expected, nAb and RBD IgG titers increased between study days 0 and 7 (**Fig. S4D-E**).

**Figure 4.**
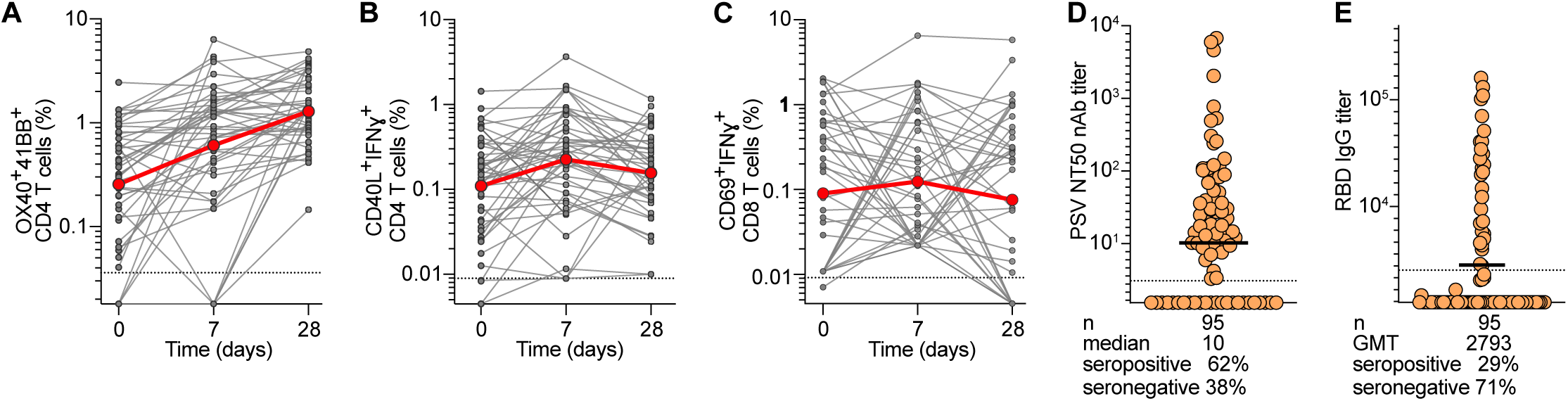
Longitudinal SARS2-specific T cell responses and Ab responses to primary SARS2 infection and acute COVID-19. **A-C.** Longitudinal combined (Spike plus non-Spike) (**A**) CD4 AIM (**B**) CD4 AIM+ICS, (**C**) CD8 IFNγ^+^ T cell responses in placebo group (n = 49) participants at study days 0, 7, and 28. Red dots and lines represent median. Dotted line = LOQ. **D.** Study day 0 pre-treatment nAb titers for all participants. Dotted black line indicates LOD; seropositivity defined by values >LOD. Bar is median. **E.** Study day 0 pre-treatment RBD IgG binding titers for all participants. Dotted black line indicates cut off for seropositivity. Bar is geometric mean titer (GMT).

To evaluate potential protective activities of early T cell responses, associations were examined between study day 0 NP viral RNA levels and the magnitude and diversity of SARS2–specific CD4 and CD8 T cell responses. The presence and magnitude of early SARS2-specific CD4 T cell responses were associated with lower levels of SARS2 NP RNA based on all three main CD4 T cell measurements of activation (OX40^+^41BB^+^ AIM Spearman r = -0.51, p = 2.6x10^-7^, **Fig. 5A**; OX40^+^CD40L^+^ AIM r = -0.45, p = 9.3X10^-6^, **Fig. S5A**) and IFNγ production (CD40L^+^IFNγ^+^, r = -0.36, p = 5x10^-4^, **Fig. 5B**). Early SARS2-specific CD8 T cell response magnitude and IFNγ production were also associated with lower levels of SARS2 NP RNA (IFNγ^+^ r = -0.44, p = 1.1x10^-5^, **Fig. 5C**; CD69^+^41BB^+^ AIM r = -0.39, p = 1.4x10^-4^, **Fig. S5C**). SARS2-specific T cell and SARS2 NP RNA associations were strongest when using combined S plus non-S T cell response measurements. Statistical associations were similar when S and non-S antigen-specific responses were evaluated separately, though generally stronger for non-S than S-specific responses (**Fig. S5D-J**). A positive correlation was observed between total SARS2-specific CD4 and CD8 T cell responses (**Fig. S5K**). A positive correlation was also observed between day 0 nAb and RBD IgG titers (**Fig. S5L**), as expected^48^. Notably, SARS2 nAb titers measured at study day 0 were also associated with lower levels of SARS2 NP RNA, but the association with nAb was weaker than the association with either day 0 SARS2-specific CD4 or CD8 T cells (r = -0.33, p = 1.2x10^-3^, **Fig. 5D**).

**Figure 5.**
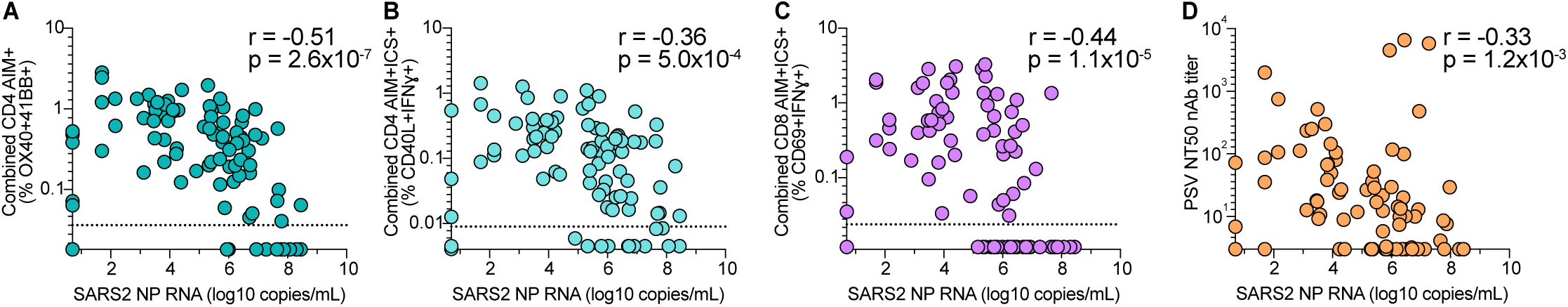
Correlative relationships between SARS2-specific adaptive immunity and upper airway viral RNA in acute COVID-19. **A-D.** Relationships between study day 0 SARS2 NP RNA and combined day 0 SARS2-specific T cell responses by (**A**) CD4 AIM^+^, (**B**) CD4 IFNγ, (**C**) CD8 IFNγ^+^ and (**D**) nAb titers.

Given the biological dependence of the antibody response on CD4 T cell help mediated by T_FH_ cells (51, 52), some statistical association between antibodies and viral load would always be expected, even if, hypothetically, viral control was exclusively mediated by T cells. Early S-specific cT_FH_ cells had relatively modest associations with SARS2 NP RNA that were weaker than total S-specific CD4 T cell AIM response associations but similar to nAb titer associations (Spearman r = -0.37 to 0.39, p = 1.6 to 2.8 x10^-4^, **Fig. S5I-J**). Analysis of covariance (ANCOVA) testing was performed to better evaluate if the putative protective relationships observed between early SARS2-specific CD4 T cell responses and SARS2 NP RNA remained after accounting for antibodies, based on day 0 nAb serostatus. Associations between total SARS2-specific CD4 or SARS2-specific cT_FH_ T cell responses and lower SARS2 NP RNA did not show a dependence on SARS2 nAb seropositivity (**Fig. S5M-O**). SARS2-specific CD8 T cell associations with lower SARS2 NP RNA was also independent of nAb seropositivity (**Fig. S5M**). Thus, early SARS2–specific CD4 and CD8 T cell responses were specifically associated with reduced upper airway SARS2 viral loads in unvaccinated, non-hospitalized cases of acute COVID-19.

Examining SARS2 antigen-specific CD4 and CD8 cytokine production and associations with viral NP RNA at study day 0 revealed that in addition to IFNγ production, production of the Th1 cytokines GzmB, IL-2, and TNF similarly correlated with lower NP viral RNA levels (Fig. S6A-F). These associations were similar for cytokine-producing CD4 and CD8 T cells recognizing both Spike and non-Spike epitopes. SARS2-specific CD4 T cells, associations were similar regardless of which cytokine the cells were producing, but for SARS2-specific CD8 T cells production of both IFNγ and GzmB was most strongly associated with decreased SARS2 NP RNA (Fig. S6D). Thus, diverse and polyfunctional SARS2-specific CD4 and CD8 T cell responses, including CD4 and CD8 T cells with profiles of cytotoxicity, may play a role in viral clearance during acute SARS-CoV-2 primary infection.

Participants in the study enrolled at a range of days post-symptom onset (PSO), which provides richness to the dataset, but raised the possibility that the immune response associations with reduced NP viral RNA levels might only be surrogates for time from infection. To test this possibility, an analysis was performed to try to account for differences in time from symptom onset to study entry, using simple log-linear regression models of the raw data to extrapolate SARS2-specific immune response and NP RNA data to day 6 PSO (the median number of days PSO for all participants) (**Fig. S7A-E**, see Methods). If the original SARS2-specific T cell associations with reduction of SARS2 NP RNA levels were only surrogates of time from infection (measured as days PSO), T cell associations should be lost upon adjustment for days PSO. If instead the T cell associations reflected a functional relationship, statistical associations between T cell responses and lower viral RNA would be expected to remain after adjustment for days PSO. Although weaker, the relationships between CD4 and CD8 T cell responses and viral NP RNA levels adjusted to 6 days PSO remained statistically significant (CD4 AIM r = -0.39, p = 1.9x10^-4^ **Fig. S7F**; CD4 IFNγ^+^ r = -0.35, p = 1.0x10^-3^ **Fig. S7G**; CD8 IFNγ^+^ r = -0.31, p = 4.4x10^-3^ **Fig. S7H**), consistent with a functional role for T cells in control of viral NP RNA levels. Associations between nAb titers and viral NP RNA levels also remained after adjustment (r = -0.32, p = 3.0x10^-3^ **Fig. S7I**). Altogether, these data are evidence of functional relationships in humans between the development of early antiviral CD4 and CD8 T cell responses, and antibodies, and control of acute SARS2 infection in the upper airway.

In summary, this study provides new insights into the relationship between acute adaptive immune responses to SARS2 and viral control. It has been hypothesized that SARS2–specific antiviral T cell responses may contribute to protection from symptomatic and severe disease by reducing viral burden and the enhancing speed of viral clearance; however, few human studies have directly measured the role of cellular immunity on viral control of primary infection (36, 40). Here, using samples from 95 individuals who participated in a randomized, controlled clinical trial, the largest virus-specific acute immune response data set of its kind was generated for adults up to 86 years of age, including many subjects at risk for progression to severe COVID-19.

## Discussion

The data demonstrate that (i) SARS2-specific CD4 and CD8 T cells could be detected in the majority of adults with outpatient COVID-19 within days from symptom onset, (ii) the T cells were polyfunctional, (iii) stronger early SARS2-specific CD4 T cell responses were associated with lower viral RNA levels in the upper airway, (iv) stronger SARS2-specific CD8 T cell responses were associated with lower viral RNA levels in the upper airway, (v) the associations of T cell responses with lower viral RNA were independent of neutralizing antibody responses and (vi) remained significant after accounting for time from COVID-19 symptom onset. The conclusions from these analyses of T cell associations with viral control were reproducible by multiple different measurements of CD4 T cell and CD8 T cell functionalities and SARS2 antigen specificities. Overall, these findings suggest that SARS2–specific T cells play a role in protective immunity to SARS2 during acute infection.

In a smaller study (n=25 acute infections sampled), Eser *et al.* found a strong association between nucleocapsid-specific IFNγ^+^ CD4 T cells and lower viral loads (40). Those results are consistent with the CD4 T cell findings here, except the findings here were that T cell responses to S or non-S antigens were both associated with reduced viral loads and were approximately equally associated with reduced viral loads though somewhat higher for non-S responses. Eser *et al.* observed less of an association for nucleocapsid-specific IFNγ^+^ CD8 T cell responses and lower viral loads, perhaps because the study sample size was limited and fewer participants developed early circulating nucleocapsid-specific IFNγ^+^ CD8 than CD4 T cell responses (40).

In a human controlled SARS2 infection study of healthy young adults (n=18 infected. Mean age of 22 years old) with a limited infectious dose (53% of individuals were infected), total CD38^+^Ki67^+^ CD8 T cells in peripheral blood were the strongest association with accelerated viral clearance (36). Those data are consistent with the CD8 T cell findings here regarding early SARS2-specific CD8 T cell activation being associated with reduced viral NP RNA. A strength of that study was the precisely defined time of infection and longitudinal tracking of viral loads and T cell responses in blood based on markers. Several factors distinguish the study reported herein. Early SARS2 Spike-and non-Spike-specific CD8 T cells both correlated with viral clearance, including when quantified by two different T cell assays (AIM and/or IFNγ production). Additionally, early SARS2 total, S-and non-S-specific CD4 T SARS2-specific responses also correlated. Furthermore, SARS2-specific cT_FH_ were also associated with lower viral loads herein. The cohort size here is substantially larger (95 compared to 18) and presumably includes subjects infected by a range of viral exposure doses. The virus-specific T cell assays herein measured CD4 and CD8 T cell epitopes spanning the Spike protein and non-S antigens including both structural and non-structural proteins and we were able to detect early circulating SARS2-specific T cells. Given that non-structural proteins are expressed earlier in infection and are better conserved across human coronaviruses, it is possible that the inclusion of non-structural protein epitopes in our peptide megapools allows for earlier and more comprehensive detection of both cross-reactive and SARS2-specific circulating T cell responses than peptide pools containing only SARS-CoV-2 structural protein epitopes (38, 53, 54). Additionally, the data herein demonstrate that the early SARS2-specific CD4 and CD8 T cells were polyfunctional, not only defined by IFNγ production (36, 40). Lastly, in the controlled infection study (36) all subjects were young healthy adults (mean age 22 years old), while the subjects herein were from a broad age range (18-86 years old) and half had comorbid conditions considered to be high risk for development of severe COVID-19. Thus, the data herein represent a broad age spectrum and are also particularly relevant for those age groups and demographics most at risk for worse outcomes due to COVID-19 like hospitalization or death. This study in no way excludes a complementary role for antibodies in control of SARS2 in unvaccinated individuals. Our study has limitations. It is intrinsically challenging to disentangle roles of T cells from roles of antibodies in combatting SARS2 in humans (15, 17, 18). This study provides evidence that SARS2-specific T cell responses exhibit selective correlates with viral clearance, but it is entirely plausible that coordinated adaptive immunity of T cells and antibodies are important in protection (14, 16, 39). Additionally, adaptive immune responses at the site of initial infection were not directly assessed in this study and may not be adequately captured by evaluation of circulating immune cells and antibodies from peripheral blood samples. Thus, a role for local, upper airway cellular and humoral immunity in protection from SARS2 cannot be excluded (16, 36, 55–57). The study was also underpowered to distinguish the myriad roles that different CD4 and CD8 T cell subsets may individually play in viral clearance, or to determine whether SARS2-specific CD4 and CD8 T cell responses are independent. We were able to demonstrate that associations between early SARS2-specific CD4 and CD8 T cell responses and NP viral RNA were independent of both nAb serostatus and time from SARS2 infection using relatively simple statistical models. While this is in agreement with other studies demonstrating a role for T cells protection against SARS2 infection (36, 39, 40), the use of more sophisticated mathematical models applied across multiple data sets may provide additional insight. Lastly, our findings focused on primary infection with ancestral SARS2 and may not be reflective of all individuals with acute COVID-19, including primary infection by SARS2 variants or SARS2 breakthrough infections.

While T cell kinetics and phenotypes in response to SARS2 infection and COVID-19 vaccination may not be identical and are shaped by exposure history (28, 41, 58), the cumulative data now indicate that SARS2-specific T cells appear to play a protective role in both primary SARS2 infection and breakthrough infections. It has been observed that in COVID-19 vaccinated individuals who experience breakthrough infections, recall responses include rapid activation of memory S-specific CD4 and CD8 T cells within the first week, which precede humoral responses, and that S-specific CD8 T cells are associated accelerated viral clearance (41, 42). Given the overall findings in the field, next generation SARS2 or pan-sarbecovirus vaccines may benefit from eliciting enhanced T cell responses, and from clinical trials that assess both baseline and vaccination-induced SARS2-specific CD4 and CD8 T cell responses and functionality.

## METHODS

### Sex as a biological variable

Our study examined similar numbers of males and females as SARS-CoV-2 infects both males and females (**Fig. 1A**). Similar findings are reported for both males and females.

### Study population and trial

ACTIV-2/A5401 was a multicenter phase 2/3 randomized controlled trial designed to evaluate the safety and antiviral and clinical efficacy of therapeutics for acute COVID-19 in non-hospitalized adults. Individuals in this study represent a subset of participants from ACTIV-2/A5401 who received a single intravenous dose of 700 mg of bamlanivimab or normal saline placebo (comparator control group) and had available clinical data (e.g., demographics, days post-symptom onset, risk for severe COVID-19; **Fig. 1A** and ref(21, 47)) and blood samples for immunologic testing. Samples from nearly half of the overall bamlanivimab 700 mg and comparator placebo groups were available for this study. Participants were from the United States and enrolled in ACTIV-2/A5401 between October-November 2020. At study entry, COVID-19 vaccines were not available to the general population in the United States and SARS2 infections could be attributed to ancestral SARS2 virus. The subjects in this study were comparable at study entry across treatment groups (**Fig. 1A** and ref(21, 47)). Inclusion criteria for ACTIV-2/A5401 included adults 18 years or older with no more than 10 days of COVID-19 symptoms and documented SARS2 infection by FDA-authorized antigen or molecular testing within seven days prior to study entry. Subjects were assigned to the bamlanivimab or placebo groups at a 1:1 ratio, and randomization was stratified by time from symptom onset (less than or ≥5 days post-symptom onset) and risk of progression to severe COVID-19 based on age (less than or ≥ 55 years old) and the presence or absence of pre-defined comorbid medical conditions (e.g., body mass index > 35kg/m^2^, hypertension, diabetes) (47). Additional information for ACTIV-2/A5401 is available at ClinicalTrials.gov (Identifier: NCT04518410) and in the primary outcomes manuscript (47). Three individuals were noted to have high NP viral loads despite high nAb titers at study day 0 (**Fig. 5D**). Notably, one of these three individuals was in the placebo group. It was subsequently re-confirmed that blood was drawn for all participants prior to bamlanivimab or placebo infusion, and that none of the placebo group participants received bamlanivimab or any other active drug treatment for COVID-19.

Partial data for day 0 RBD IgG titers by group, similar to the data shown in **Fig. S4C** but with smaller sample sizes and without using a strict mean fluorescence intensity (MFI)-based cutoff, was previously published (21). Although combined day 28 AIM and AIM+ICS data was published to show T cell response rates between the treatment and placebo groups (21), it was not presented as longitudinal data (as shown in Fig. **4A-C**, **S4A**). Day 0 and 7 T cell data have not been previously published.

### SARS2 viral RNA

Samples were collected, processed, and analyzed as previously reported (47). In brief, nasopharyngeal swab samples were collected by ACTIV-2/A5401 trial staff at designated study sites on assigned study days (days 0, 7, 14, and 28) using standardized swabs and validated collection and storage procedures. nasopharyngeal swabs were frozen upon collection and stored at −80 °C (−65 °C to −95 °C) until shipped on dry ice to a central laboratory (University of Washington) for extraction, amplification and quantitative detection testing using validated, previously published methods using the Abbott m2000sp/rt system with a validated controls and standards for correlation with cycle threshold (Ct) and viral load. The limit of detection (LOD) was 1.4 log10 copies/mL, lower limit of quantification (LLOQ) was 2 log10 copies/mL, and upper limit of quantification (ULOQ) was 7 log10 copies/mL for this assay. For samples with viral RNA levels above the ULOQ, samples were diluted, and the assay repeated to obtain a quantitative value.

### PBMC and viability-based quality control

Peripheral blood was collected by ACTIV-2/A5401 trial staff at designated study sites on assigned study days. Serum and peripheral blood mononuclear cells (PBMC) for immunologic testing were isolated from whole blood using standard operating procedures. PBMC were cryopreserved and stored in liquid nitrogen until ready for use, then thawed at 37°C, resuspended in warm complete RPMI medium with 5% human AB serum (Gemini Bioproducts) and benzonase and centrifuged to remove cryopreservation medium. Following washing, cell counts were performed, and viability assessed using the Muse Count & Viability Kit (Muse Cell Analyzer; Luminex). A PBMC viability threshold was set at ≥ 75% for all samples for inclusion in data analysis. PBMC with viability below this threshold failed to appropriately respond to control stimuli. Four of 95 PBMC samples failed to meet the viability cutoff and were excluded from analyses. PBMC were resuspended to achieve a final concentration of 100,000 PBMC/100 μL for plating in 96-well format for T cell assays. Cryopreserved convalescent COVID-19 donor PBMC from known positive responders obtained from the Sette lab under an LJI IRB-approved protocol (VD-214) served as batch controls across T cell assays. These PBMC were handled in the same fashion as ACTIV-2/A5401 PBMC samples. Additional quality control metrics were also applied, as described below for the individual T cell assays.

### Activation induced marker assay (AIM)

As previously described (21), PBMC were cultured in 96-well U-bottom plates for 24 hours at 37°C in an incubator with 5% CO2 in the presence of a single stimulus per well: a negative control (equimolar amount of DMSO vehicle), a positive control (Staphylococcal enterotoxin B at 1 μg/mL), or a single SARS2 MP (1 μg/mL per MP) (20, 28, 59) containing Spike (S) (20, 59) or paired SARS2 non-Spike epitope MPs (CD4-RE or CD8-RE dominant and subdominant MPs (28)). The Spike MP was comprised of overlapping peptides spanning the full-length ancestral Spike protein. The CD4-RE pools (“non-Spike”) consisted of experimentally validated and optimized class II epitopes from the remainder of the SARS2 proteome (outside of the Spike open reading frame (ORF)). CD8 AIM responses were calculated using the AIM+ICS assay as PBMC counts were limited and did not allow for CD8-RE MP wells to be plated for both AIM and AIM+ICS assays (see AIM+ICS assay section below for additional experimental details).

PBMC were plated at 1 x 10^6^ PBMC per MP stimulation well and between 0.5-1 x 10^6^ PBMC per control well; negative controls were plated in duplicate. Prior to stimulation, PBMC were incubated at 37°C for 15 minutes with 0.5 μg/mL anti-human CD40 blocking antibody (Miltenyi Biotec) per well. Chemokine receptor antibodies were also added to each well on day 1 (see ref 21 for antibodies used in the AIM assay). After a 24-hour incubation, the plates were centrifuged, cells washed with PBS, then stained with LIVE/DEAD Fixable Blue (Invitrogen) 1:1000 in PBS with Fc block (5 μL/sample; BD Biosciences (BD)) for 15 minutes at room temperature, washed with FACS buffer (3% FBS in DPBS without calcium or magnesium), surface stained (see ref 21 for surface staining panel; 30 minutes at 4°C), fixed with BD Cytofix Fixation Buffer (4°C for 20 minutes), and stored at 4°C (up to overnight) until flow cytometric analysis was performed using a 5-laser Cytek Aurora (Cytek Biosciences). Flow cytometric data was acquired separately for each sample and stimulation condition; no samples were pooled. Analysis was performed using FlowJo (BD), and AIM^+^ gates were drawn based on MP-stimulated responses relative to negative (DMSO) and positive (SEB) controls. T cell assays on two SARS2 convalescent control PBMC samples were run with each set of experimental PBMC samples to account for batch effects; aliquots from the same PBMC controls were used across batches. A single gating strategy was applied to all samples across all batches unless there was a clear batch-specific effect to justify modifying the gates for a specific batch of samples. Our standard acceptable background (% AIM^+^ T cells) for DMSO gates is 0.1. DMSO gates were set to target average background values ≤ 0.1 using the same flow cytometry gating strategy for all samples across all batches. CD4 and CD8 T cell responses to MP versus DMSO stimulation are shown in Fig. S2K-M and S3G-H. Higher background was allowed for CD8 AIM (CD69^+^41BB^+^) assays (Fig. S3G) based on higher background observed for CD8 AIM both here and in prior studies (21). ICS (CD69^+^IFNγ^+^) was preferentially used as the main readout for SARS2-specific CD8 T cell responses given lower background (Fig. S3H) with comparable response rates to CD8 AIM.

PBMC quality was evaluated by measuring the median response to SEB for all samples, and samples with responses <50% of the overall median SEB response were excluded from downstream analyses. Two DMSO negative control well replicates were run for each sample and the average of the individual DMSO wells for each sample was calculated. T cell responses were verified by stimulation index (SI) as a quality control. SI was calculated for each sample as the fold change in responses to MP-stimulation versus the average response to DMSO for the same subject (as shown in **Fig. S2N-P** and **Fig. S3I-J**). An SI cutoff of 2 was used for determining AIM^+^ CD4 T cell responses and 3 for AIM^+^ CD8 T cell responses. The higher SI cutoff for CD8 T cell responses was applied based on higher background for CD8 AIM assays. When a non-zero DMSO value was needed for calculating the stimulation index, the minimum DMSO signal was set to 0.005%. For each MP-stimulated sample, the average DMSO value was subtracted from the MP response to calculate the background subtracted MP response. Background subtracted S and non-S values are shown for all figures. Combined AIM responses were calculated as the mathematical sum of the background subtracted responses to individual (S plus non-S) MPs. In the case of negative AIM responses to both S and non-S megapools, or summative responses that were less than the lower limit of quantitation (LOQ) for the assay, as described below, the combined AIM response was also considered negative. The geometric mean of all DMSO control wells for all samples for each assay was calculated to determine the LOQ for the assay. Positive responders were defined as those with background subtracted responses greater than the LOQ. The baseline for the y-axis was set to 0.5 x LOQ. All non-responders were set to this baseline.

### Hybrid activation induced marker plus intracellular cytokine staining assay (AIM+ICS)

AIM+ICS assays were also performed as previously described (21). For CD4 T cells, intracellular cytokine expression was measured in conjunction with surface CD40L. For CD8 T cells, intracellular cytokine staining was measured in conjunction with surface CD69. PBMC were thawed and plated in parallel and as AIM assays but with the addition of wells for CD8-RE MP (paired dominant and non-dominant epitopes (28)) stimulation. The CD8-RE pools (“non-Spike”) consisted of optimized, experimentally validated class I epitopes from the remainder of the SARS2 proteome (outside of the Spike ORF) (28). Cells were incubated with anti-CD40 blocking antibody as for AIM assays, but no chemokine receptor antibodies were added on day 1 except for CXCR5 for identification of circulating T_FH_ cells (see ref 21 for antibodies used). After 20-22 hours, PMA (0.05 μg/mL) and ionomycin (0.25 μg/mL) were added to the positive (ICS) control wells. Two hours later, 0.25 μL/well of GolgiStop (BD) and GolgiPlug (BD) and the AIM marker antibodies (see ref(21)) were added to all samples, and the plates were incubated for another 4 hours at 37°C (in a 5% CO_2_ incubator). Cells were then washed, surface stained for 30 minutes at 4°C, fixed and washed using Cytofix/Cytoperm (BD) per the manufacturers’ protocol. Intracellular cytokine staining was then performed using antibodies diluted in Perm/Wash Buffer (BD) for 30 minutes at 4°C. Cells were washed with Stain Buffer with FCS (BD) and stored in this buffer at 4°C until flow cytometric analysis was performed using a Cytek Aurora. Flow cytometric data was acquired separately for each sample and stimulation condition.

Gating and quality control evaluations with SEB responses and SI calculations were performed as described above for AIM assays. PMA + ionomycin served as an additional control for cytokine production (ICS). For each MP-stimulated sample, the average DMSO value was subtracted from the MP response to calculate the background subtracted MP response. Background subtracted values are shown for all figures unless indicated. Combined responses were calculated as the sum of the background subtracted responses to individual MPs (S and CD4-RE for CD4 T cells, S and CD8-RE for CD8 T cells). In the case of negative ICS responses to both S and non-S MPs, the combined response was also considered negative. The LOQ and baseline was calculated and plotted as described above for AIM assays.

### Binding antibody titers

Serum SARS2-specific binding antibody assays were performed using the Bio-Plex Pro Human SARS2 IgG 4-Plex Panel serology assay (Bio-Rad 12014634) according to the manufacturer’s protocol, as previously described (21). MFI-based cutoffs for seropositivity are indicated in the figures.

### Pseudovirus neutralizing antibody titers

Half-maximal neutralizing antibody titers (NT50) were generated using a lentiviral pseudovirus assay, as previously described (60). Briefly, PSV was generated in HEK293T cells co-transfected with SARS2 Spike expression and pNL4–3.luc.R-E-mCherry-luciferase reporter packaging plasmids. Serum was isolated from the peripheral blood of study subjects using standard protocols, frozen and shipped to the testing site without thawing until ready for use. The serum was then serially diluted and incubated with SARS2 PSV, then HEK293T cells stably expressing human angiotensin converting enzyme 2. Serum dilutions were run in triplicate. The percent inhibition was calculated using relative light units emitted by the reporter at each dilution and plotted versus the log serum dilution. The NT50 was then derived by non-linear regression. The limit of detection for the assay was defined by the lowest dilution tested (1:3) as indicated in figures.

### Statistics

Statistical analyses were performed in GraphPad Prism 10 (GraphPad Software) and Microsoft Excel. Comparisons of SARS2-specific T cell response magnitudes and antibody titers between treatment and placebo groups for equivalent conditions were determined by Mann-Whitney tests. Fisher’s exact tests were used to compare participant characteristics (**Fig. 1A**) and to compare SARS2-specific T cell positive and negative response rates between treatment and control groups. Kruskal-Wallis nonparametric tests with post-hoc Dunn’s multiple comparison tests were used for assessing T cell responses across more than one stimulation condition or cell type. Relationships between SARS2 nasopharyngeal RNA levels and immune responses were assessed using nonparametric Spearman correlations. One-way ANCOVA were performed on unadjusted viral RNA and immune response data to determine if the relationships between SARS2 NP RNA levels and SARS2-specific T cell responses at study day 0 were dependent on SARS2 nAb serostatus (**Fig. S5M-O**).

Individual variation in days from symptom onset to study entry (day 0) was accounted for using a log-linear regression model (**Fig. S7A-E**). Adjusted SARS2 NP RNA levels (**Fig. S7A**) and adaptive immune responses (log_10_ transformed CD4 T cell responses, CD8 T cell responses, and nAb titers; **Fig. S7B-E**) at 6 days post-symptom onset (the median time from symptom onset at study entry) were calculated for all participants using the linear regression models shown in **Fig. S7A-E** to adjust the data shown in **Fig. 5A-D**. Data from five individuals were excluded from this analysis as outliers as these individuals had undetectable viral RNA on study day 0 but were reportedly 4-5 days PSO as study entry. This included four values from individuals who consistently tested negative for viral RNA (from both nasal and NP swabs) on all study days, and a value for one individual who tested negative for viral NP RNA on study day 0 but who subsequently tested positive for viral NP RNA on study days 3, 7, and 14. The adjusted NP viral RNA and adaptive immunity data was plotted and analyzed using nonparametric Spearman correlations to be able to compare the original (unadjusted) data (Fig. 5A-D) to the adjusted data (**Fig. S7F-I**) using the same correlation method.

For all analyses, a two-sided 5% type I error rate was used, without adjustment for multiple comparisons, except as noted above. Additional details can be found in the results, figures and corresponding legends.

### Study approval

The ACTIV-2/A5401 clinical trial protocol (ClinicalTrials.gov Identifier: NCT04518410) was approved by a central institutional review board (IRB); Advarra (Pro00045266). The La Jolla Institute for Immunology (LJI) IRB provided additional oversight for this study. All individuals enrolled in ACTIV-2/A5401 provided written informed consent for participation.

## Data availability

The authors confirm that the source data underlying the findings are fully available. Due to ethical restrictions, additional ACTIV-2/A5401 clinical trial study data beyond what is presented in this manuscript and supplement are available upon request from the AIDS Clinical Trials Group (ACTG) and with the written agreement of ACTG and the manufacturer of the investigational product. Completion of a data use agreement may be required. Values for all data points in graphs are reported in the Supporting Data Values file.

## AUTHOR CONTRIBUTIONS

Conceptualization, S.I.R., D.M.S., and S.C.; Investigation, S.I.R., P.G.L., F.F., A.H.; Formal Analysis, S.I.R., B.P, S.C.; Patient Recruitment and Samples D.M.S.; Material Resources, A.G., D.W., P.K., A.S.; Writing, S.I.R., F.F., D.M.S. and S.C.; Supervision D.M.S. and S.C.

## ACKNOWLEDGMENTS

We would like to thank the study participants, site staff and investigators, members of the study team and community advisory board, the AIDS Clinical Trials Group (ACTG), the Harvard Center for Biostatistics in AIDS Research (CBAR) and ACTG Statistical and Data Analysis Center (SDAC), the National Institute of Allergy and Infectious Diseases (NIAID) / Division of AIDS (DAIDS), Bill Erhardt, the Accelerating COVID-19 Therapeutic Interventions and Vaccines (ACTIV) partnership, and PPD for making this study possible. See Supplemental Acknowledgments for additional details.

