## Supplementary Figures 1-7 for "Early antiviral CD4 and CD8 T cell responses and antibodies are associated with upper respiratory tract clearance of SARS-CoV-2"

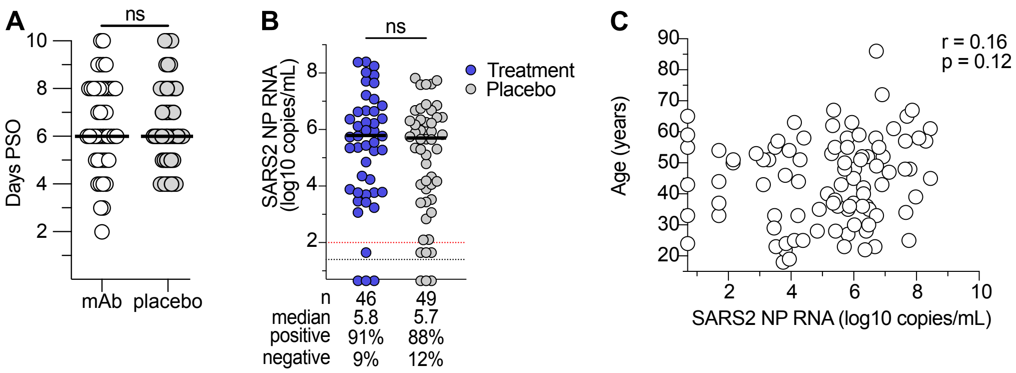


**Figure S1. Days from symptom onset to study entry and SARS2 NP RNA at study entry by clinical trial group.**
**A.** Days PSO at study day 0 for bamlanivimab (mAb; n = 46) and placebo group participants (n = 49); line = median (day 6). **B.** SARS2 NP RNA at day 0 by treatment group (Treatment = bamlanivimab). **C.** Relationship between day 0 SARS2 NP RNA and participant age. Lines and bars as in **Fig. 1B-C**. ns = not significant by Mann-Whitney test. **Related to Figure 1.**

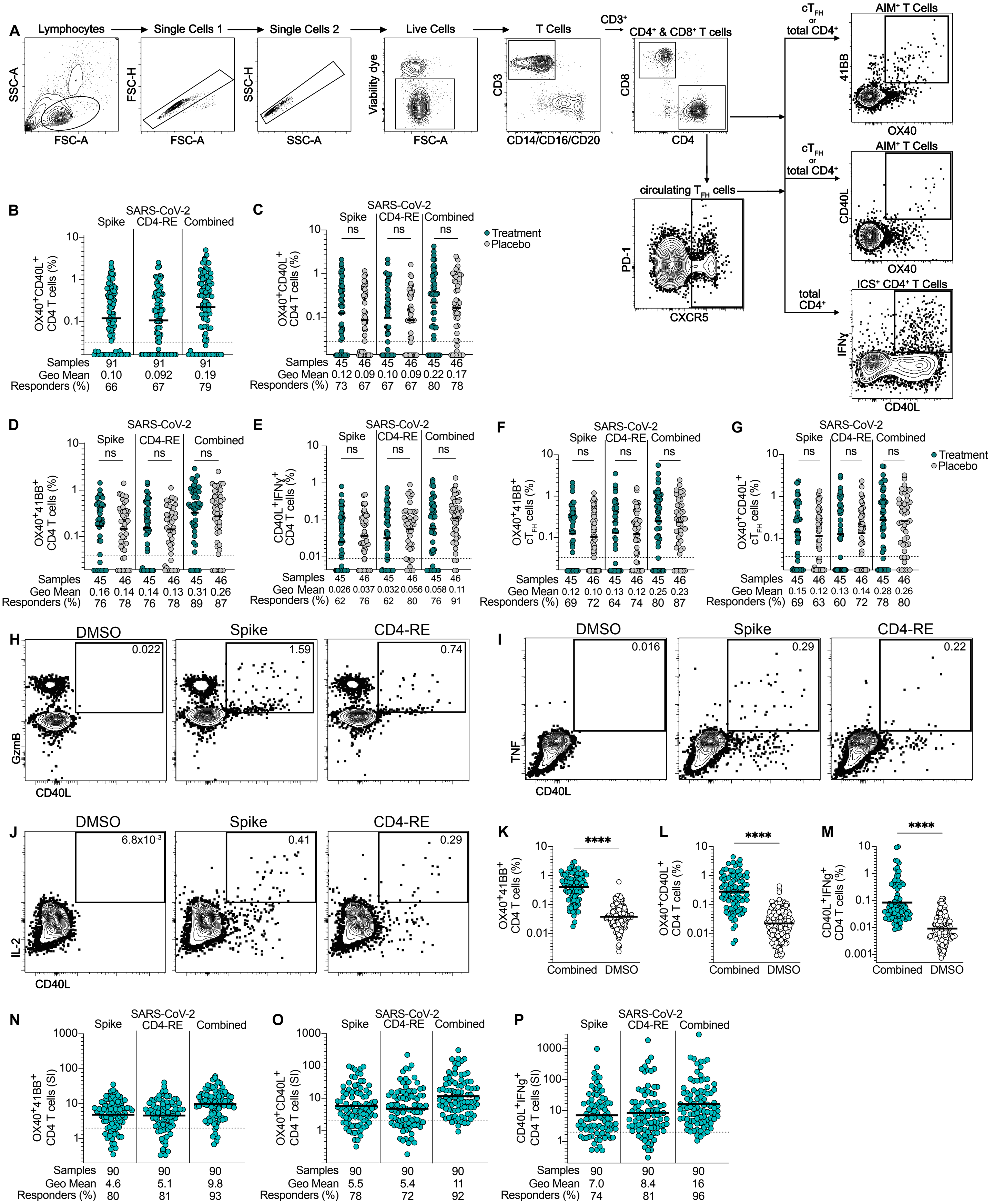

A
Lymphocytes
Single Cells 1
Single Cells 2
Live Cells
T Cells
CD3+
CD4+ & CD8+ T cells
SSC-A
FSC-A
FSC-H
SSC-H
Viability dye
CD3
CD14/CD16/CD20
CD8
CD4
cTFH or total CD4+
AIM+ T Cells
41BB
OX40
CD40L
circulating TFH cells
PD-1
CXCR5
total CD4+
ICS+ CD4+ T Cells
IFNγ
B
SARS-CoV-2
Spike
CD4-RE
Combined
OX40+CD40L+ CD4 T cells (%)
Samples 91 91 91
Geo Mean 0.10 0.092 0.19
Responders (%) 66 67 79
C
Treatment
Placebo
ns
Samples 45 46 45 46 45 46
Geo Mean 0.12 0.09 0.10 0.09 0.22 0.17
Responders (%) 73 67 67 67 80 78
D
OX40+41BB+ CD4 T cells (%)
Samples 45 46 45 46 45 46
Geo Mean 0.16 0.14 0.14 0.13 0.31 0.26
Responders (%) 76 78 76 78 89 87
E
CD40L+IFNγ+ CD4 T cells (%)
Samples 45 46 45 46 45 46
Geo Mean 0.026 0.037 0.032 0.056 0.058 0.11
Responders (%) 62 76 62 80 76 91
F
OX40+41BB+ cTFH cells (%)
Samples 45 46 45 46 45 46
Geo Mean 0.12 0.10 0.13 0.12 0.25 0.23
Responders (%) 69 72 64 74 80 87
G
OX40+CD40L+ cTFH cells (%)
Samples 45 46 45 46 45 46
Geo Mean 0.15 0.12 0.13 0.14 0.28 0.26
Responders (%) 69 63 60 72 78 80
H
DMSO
0.022
1.59
0.74
GzmB
I
0.016
0.29
0.22
TNF
J
6.8x10-3
0.41
0.29
IL-2
K
****
Combined
DMSO
L
M
CD40L+IFNg+ CD4 T cells (%)
N
OX40+41BB+ CD4 T cells (SI)
Samples 90 90 90
Geo Mean 4.6 5.1 9.8
Responders (%) 80 81 93
O
OX40+CD40L+ CD4 T cells (SI)
Samples 90 90 90
Geo Mean 5.5 5.4 11
Responders (%) 78 72 92
P
CD40L+IFNg+ CD4 T cells (SI)
Samples 90 90 90
Geo Mean 7.0 8.4 16
Responders (%) 74 81 96

**Figure S2. Antigen-specific CD4 T cell responses to primary SARS2 infection and acute COVID-19.**

**A.** Gating strategy and representative flow cytometry gating for CD4 AIM and AIM+ICS assay readouts using IFNγ as the example cytokine shown. **B.** Study day 0 Spike, CD4-RE, or Combined (Spike + CD4-RE) $AIM^+$ CD4 T cells by surface $OX40^+CD40L^+$. **C-E** Study day 0 Spike, CD4-RE, and combined total CD4 T cells by clinical trial assignment (bamlanivimab = "Treatment", saline = "Placebo") by (**C**) surface $OX40^+CD40L^+$, (**D**) $OX40^+41BB^+$, or (**E**) surface CD40L and intracellular IFNγ. **F-G.** Study day 0 Spike, CD4-RE, and combined circulating TFH cells by surface (**F**) $OX40^+41BB^+$ or (**G**) $OX40^+CD40L^+$. **H-J**. Example flow cytometry gating and frequencies for total CD4 T cells based on surface $CD40L^+$ and intracellular production of (**H**) GzmB, (**I**) TNF, (**J**) IL-2. **K-M.** % SARS2-specific of total CD4 T cells for MP versus DMSO stimulated by (**K**) $OX40^+41BB^+$ (**L**) $OX40^+CD40L^+$, and (**M**) $CD40L^+IFN\gamma^+$. **N-P.** Spike, non-Spike (CD4-RE) and total (Spike + CD4-RE = Combined) SARS2-specific CD4 T cells by (**N**) $OX40^+41BB^+$, (**O**) $OX40^+CD40L^+$, and (**P**) $CD40L^+IFN\gamma^+$ by stimulation index (SI, fold-change relative to DMSO control; see Methods for details). Bars, dotted lines, and flow cytometry gate labels as in **Fig. 2**. ns = not significant between Treatment and Placebo group by Mann-Whitney for comparison of the same stimulation condition. **** = $p \leq 0.0001$ by Mann-Whitney. **Related to Figure 2.**

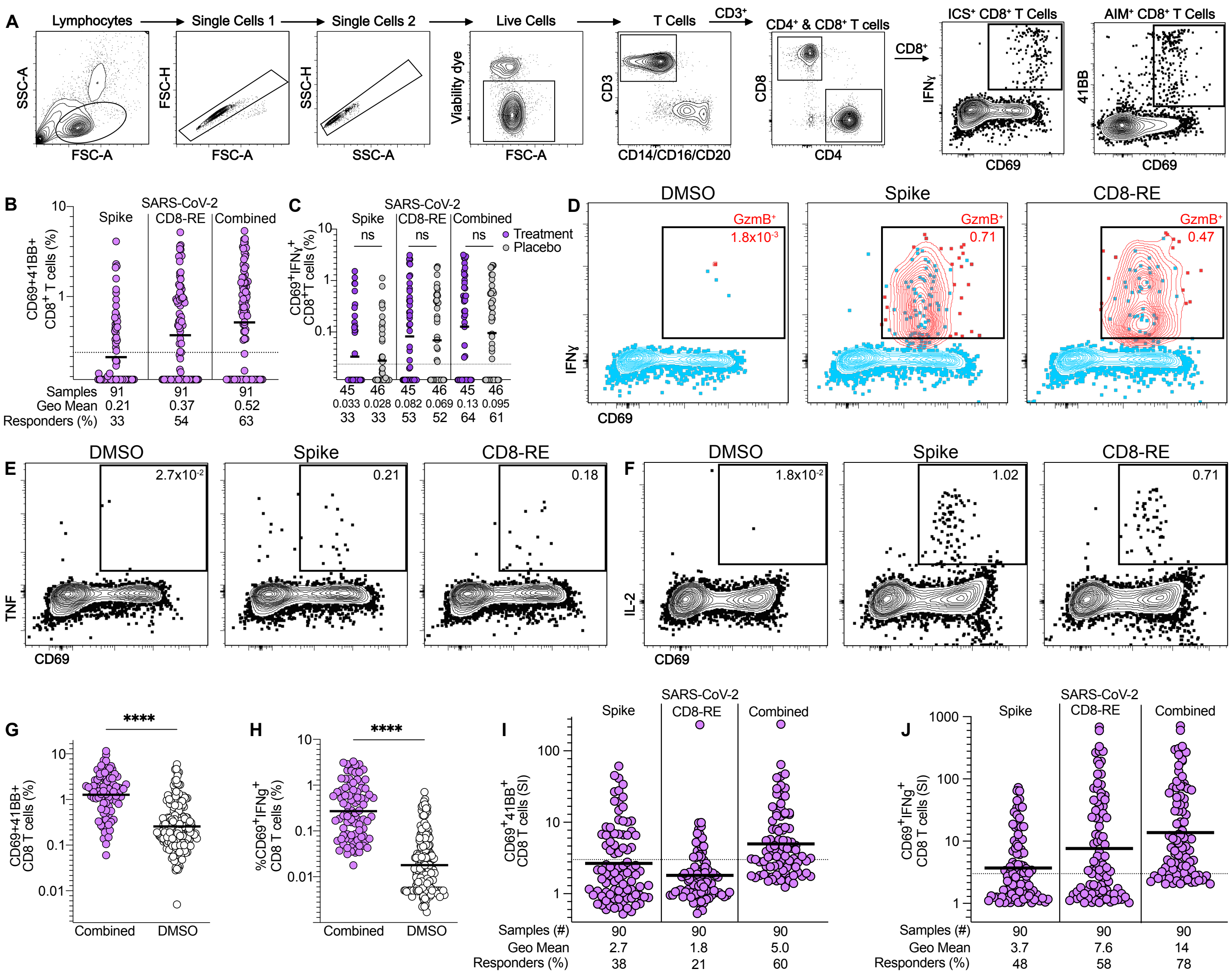


**Figure S3. Antigen-specific CD8 T cell responses to primary SARS2 infection and acute COVID-19.**

**A.** Gating strategy and representative flow cytometry gating for CD8 AIM and main ICS assay readouts. **B-C.** Study day 0 Spike, CD8-RE, or Combined CD8 responses by (**B**) AIM ($CD69^+41BB^+$) for all participants or (**C**) main ICS readout ($CD69^+IFN\gamma^+$) by clinical trial assignment (bamlanivimab = "Treatment", saline = "Placebo"). Bars and lines as in **Fig. 3**. ns = not significant between Treatment and Placebo group by Mann-Whitney for comparison of the same stimulation condition. **D-F.** Representative flow cytometry gating and frequencies for other intracellular cytokine production by SARS2-specific surface $CD69^+$ CD8 T cells for (**D**) GzmB (frequency of triple positive red cells out of total $CD69^+IFN\gamma^+$; other cells shown in light blue), (**E**) TNF, (**F**) IL-2. **G-H.** % SARS2-specific of total CD8 T cells for MP stimulated conditions versus DMSO control conditions by (**G**) $CD69^+41BB^+$ and (**H**) $CD69^+IFN\gamma^+$. **I-J.** Spike, non-Spike (CD8-RE) and total (Spike + CD8-RE = Combined) SARS2-specific CD8 T cells by (**I**) $CD69^+41BB^+$ and (**J**) $CD69^+IFN\gamma^+$ by stimulation index (SI, fold-change relative to DMSO control; see Methods). ns = not significant, **** = p ≤ 0.0001 by Mann-Whitney. **Related to Figure 3.**

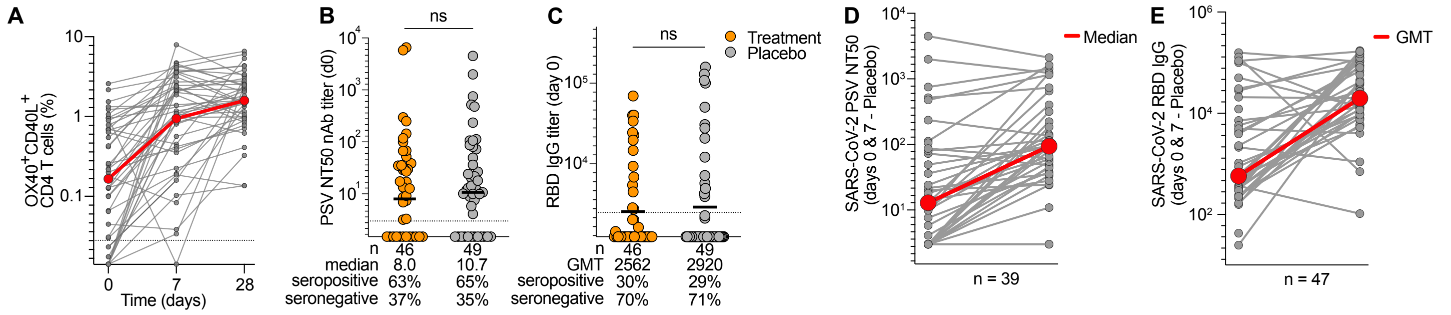


**Figure S4. Humoral responses to primary SARS2 infection and acute COVID-19.**

**A.** Longitudinal AIM$^+$ CD4 T cell responses by surface OX40$^+$CD40L$^+$ in the placebo group (n = 49) at study days 0, 7, and 28. Red line is median for each time point. **B-C.** Day 0 (**B**) nAb titers and (**C**) RBD IgG titers by clinical trial assignment (bamlanivimab = "Treatment", saline = "Placebo"). **D-E.** Day 0 and 7 (**D**) nAb titers and (**E**) RBD IgG titers for the placebo group (n = 49) participants. Bars and lines as in **Fig. 4** unless labeled differently. **Related to Figure 4.**

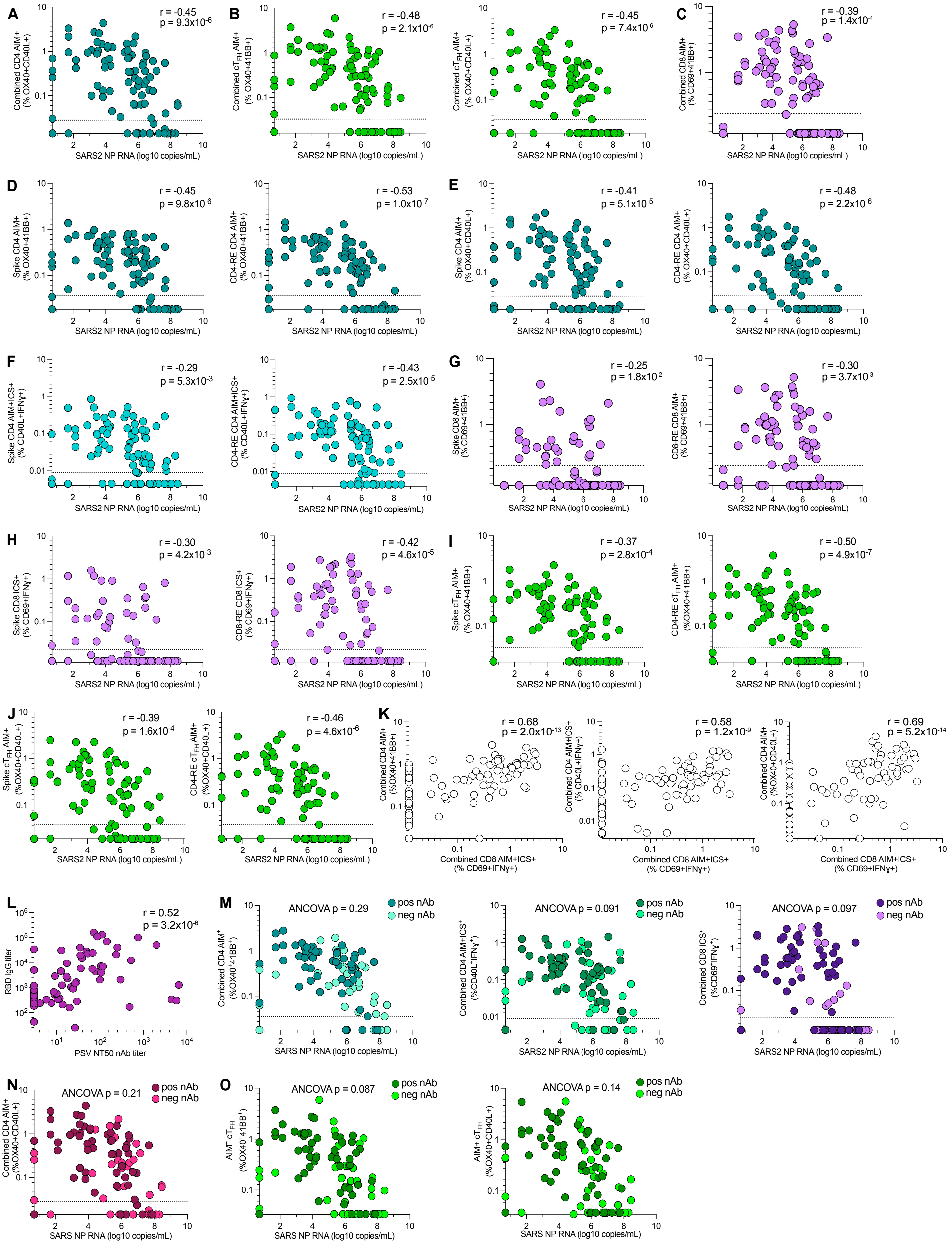
A
Combined CD4 AIM+ (% OX40+CD40L+)
r = -0.45
p = 9.3x10-6
SARS2 NP RNA (log10 copies/mL)
B
Combined cTFH AIM+ (% OX40+41BB+)
r = -0.48
p = 2.1x10-6
Combined cTFH AIM+ (% OX40+CD40L+)
r = -0.45
p = 7.4x10-6
C
Combined CD8 AIM+ (% CD69+41BB+)
r = -0.39
p = 1.4x10-4
D
Spike CD4 AIM+ (% OX40+41BB+)
r = -0.45
p = 9.8x10-6
CD4-RE CD4 AIM+ (% OX40+41BB+)
r = -0.53
p = 1.0x10-7
E
Spike CD4 AIM+ (% OX40+CD40L+)
r = -0.41
p = 5.1x10-5
CD4-RE CD4 AIM+ (% OX40+CD40L+)
r = -0.48
p = 2.2x10-6
F
Spike CD4 AIM+ICS+ (% CD40L+IFNγ+)
r = -0.29
p = 5.3x10-3
CD4-RE CD4 AIM+ICS+ (% CD40L+IFNγ+)
r = -0.43
p = 2.5x10-5
G
Spike CD8 AIM+ (% CD69+41BB+)
r = -0.25
p = 1.8x10-2
CD8-RE CD8 AIM+ (% CD69+41BB+)
r = -0.30
p = 3.7x10-3
H
Spike CD8 ICS+ (% CD69+IFNγ+)
r = -0.30
p = 4.2x10-3
CD8-RE CD8 ICS+ (% CD69+IFNγ+)
r = -0.42
p = 4.6x10-5
I
Spike cTFH AIM+ (% OX40+41BB+)
r = -0.37
p = 2.8x10-4
CD4-RE cTFH AIM+ (%OX40+41BB+)
r = -0.50
p = 4.9x10-7
J
Spike cTFH AIM+ (%OX40+CD40L+)
r = -0.39
p = 1.6x10-4
CD4-RE cTFH AIM+ (%OX40+CD40L+)
r = -0.46
p = 4.6x10-6
K
Combined CD4 AIM+ (%OX40+41BB+)
r = 0.68
p = 2.0x10-13
Combined CD4 AIM+ICS+ (% CD40L+IFNγ+)
r = 0.58
p = 1.2x10-9
Combined CD4 AIM+ (%OX40+CD40L+)
r = 0.69
p = 5.2x10-14
Combined CD8 AIM+ICS+ (% CD69+IFNγ+)
L
RBD IgG titer
r = 0.52
p = 3.2x10-6
PSV NT50 nAb titer
M
ANCOVA p = 0.29
pos nAb
neg nAb
Combined CD4 AIM+ (%OX40+41BB+)
SARS NP RNA (log10 copies/mL)
ANCOVA p = 0.091
Combined CD4 AIM+ICS+ (%CD40L+IFNγ+)
ANCOVA p = 0.097
Combined CD8 ICS+ (%CD69+IFNγ+)
N
ANCOVA p = 0.21
Combined CD4 AIM+ (%OX40+CD40L+)
O
ANCOVA p = 0.087
AIM+ cTFH (%OX40+41BB+)
ANCOVA p = 0.14
AIM+ cTFH (%OX40+CD40L+)

**Figure S5. Unadjusted correlative relationships between virus-specific immune responses and upper airway SARS2 RNA levels. A-D.** Correlation analyses similar to **Fig. 5A-C** but for study day 0 SARS2 NP RNA and SARS2-specific T cell responses by (**A**) CD4 T cell $OX40^{+}CD40L^{+}$ AIM; (**B**) circulating $T_{FH}$ by $OX40^{+}41BB^{+}$ AIM or $OX40^{+}CD40L^{+}$ AIM; (**C**) CD8 T cell $CD69^{+}41BB^{+}$ AIM. **D-J.** Relationships for study day 0 SARS2 NP RNA and SARS2-specific T cell responses as in **Fig. 5A-C** and **Fig. S5A-C** split by responses to S or non-S MP stimulation for (**D**) CD4 T cell $OX40^{+}41BB^{+}$ AIM (**E**) CD4 T cell $OX40^{+}CD40L^{+}$ AIM, (**F**) CD4 T cell $CD40L^{+}IFN\gamma^{+}$, (G) CD8 T cell $CD69^{+}41BB^{+}$ AIM, (**H**) CD8 T cell $CD69^{+}IFN\gamma^{+}$, (**I**) circulating $T_{FH}$ by $OX40^{+}41BB^{+}$ AIM, (**J**) circulating $T_{FH}$ by $OX40^{+}CD40L^{+}$ AIM. **K.** Relationships between study day 0 SARS2-specific combined CD4 and CD8 responses by AIM and/or ICS. **L.** Relationship between day 0 nAb and RBD IgG titers. **M-O.** Impact of study day 0 nAb serostatus on relationships between study day 0 T cell responses and SARS2 NP RNA shown in **Fig. S5B** (**O** for circulating $T_{FH}$), **Figs. 5A-C** and **Fig. S5A** (**M** and **N** for total CD4 and CD8) by ANCOVA. All correlations by two-tailed nonparametric testing; r = Spearman's rank correlation coefficient. **Related to Figure 5.**

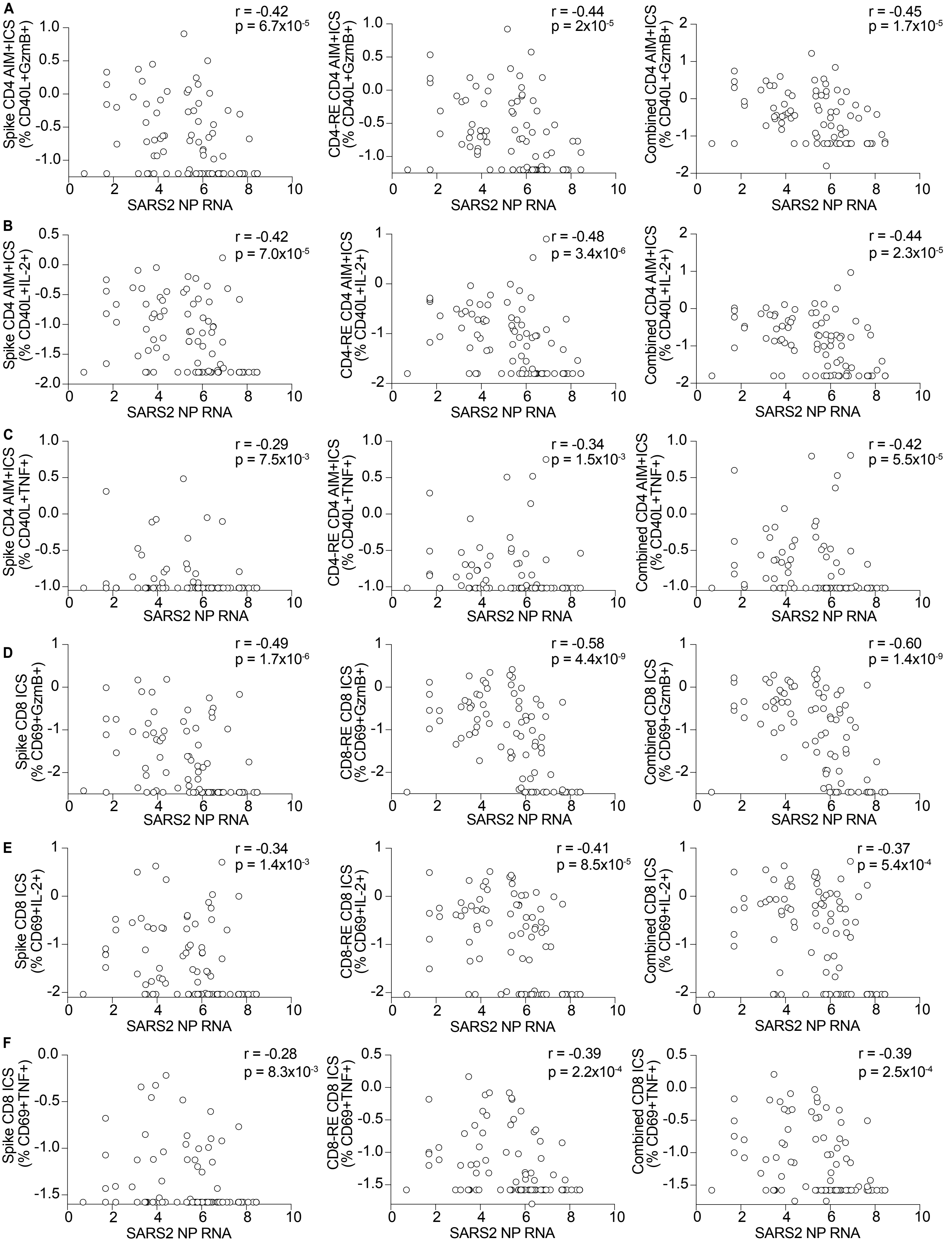
A
Spike CD4 AIM+ICS (% CD40L+GzmB+)
r = -0.42
p = 6.7x10$^{-5}$
CD4-RE CD4 AIM+ICS (% CD40L+GzmB+)
r = -0.44
p = 2x10$^{-5}$
Combined CD4 AIM+ICS (% CD40L+GzmB+)
r = -0.45
p = 1.7x10$^{-5}$
B
Spike CD4 AIM+ICS (% CD40L+IL-2+)
r = -0.42
p = 7.0x10$^{-5}$
CD4-RE CD4 AIM+ICS (% CD40L+IL-2+)
r = -0.48
p = 3.4x10$^{-6}$
Combined CD4 AIM+ICS (% CD40L+IL-2+)
r = -0.44
p = 2.3x10$^{-5}$
C
Spike CD4 AIM+ICS (% CD40L+TNF+)
r = -0.29
p = 7.5x10$^{-3}$
CD4-RE CD4 AIM+ICS (% CD40L+TNF+)
r = -0.34
p = 1.5x10$^{-3}$
Combined CD4 AIM+ICS (% CD40L+TNF+)
r = -0.42
p = 5.5x10$^{-5}$
D
Spike CD8 ICS (% CD69+GzmB+)
r = -0.49
p = 1.7x10$^{-6}$
CD8-RE CD8 ICS (% CD69+GzmB+)
r = -0.58
p = 4.4x10$^{-9}$
Combined CD8 ICS (% CD69+GzmB+)
r = -0.60
p = 1.4x10$^{-9}$
E
Spike CD8 ICS (% CD69+IL-2+)
r = -0.34
p = 1.4x10$^{-3}$
CD8-RE CD8 ICS (% CD69+IL-2+)
r = -0.41
p = 8.5x10$^{-5}$
Combined CD8 ICS (% CD69+IL-2+)
r = -0.37
p = 5.4x10$^{-4}$
F
Spike CD8 ICS (% CD69+TNF+)
r = -0.28
p = 8.3x10$^{-3}$
CD8-RE CD8 ICS (% CD69+TNF+)
r = -0.39
p = 2.2x10$^{-4}$
Combined CD8 ICS (% CD69+TNF+)
r = -0.39
p = 2.5x10$^{-4}$
SARS2 NP RNA

**Figure S6. Additional unadjusted correlative relationships between SARS2-specific T cell cytokine production and SARS2 NP RNA levels. A-C.** Correlation analyses for study day 0 SARS2 NP RNA and SARS2 Spike (left), non-Spike (CD4-RE; middle), and combined (right) CD4 T cell cytokine production by CD4 T cell AIM+ICS: (**A**) CD40L$^{+}$GzmB$^{+}$ (**B**) CD40L$^{+}$IL-2$^{+}$ (**C**) CD40L$^{+}$TNF$^{+}$. **D-F.** Correlation analyses for study day 0 SARS2 NP RNA and SARS2 Spike (left), non-Spike (CD4-RE; middle), and combined (right) CD8 T cell cytokine production by CD8 T cell ICS: (**D**) CD69$^{+}$IFNγ$^{+}$GzmB$^{+}$, (**E**) CD69$^{+}$IL-2$^{+}$, (**F**) CD69$^{+}$TNF$^{+}$. All correlations by two-tailed nonparametric testing; r = Spearman's rank correlation coefficient. **Related to Figure 5.**

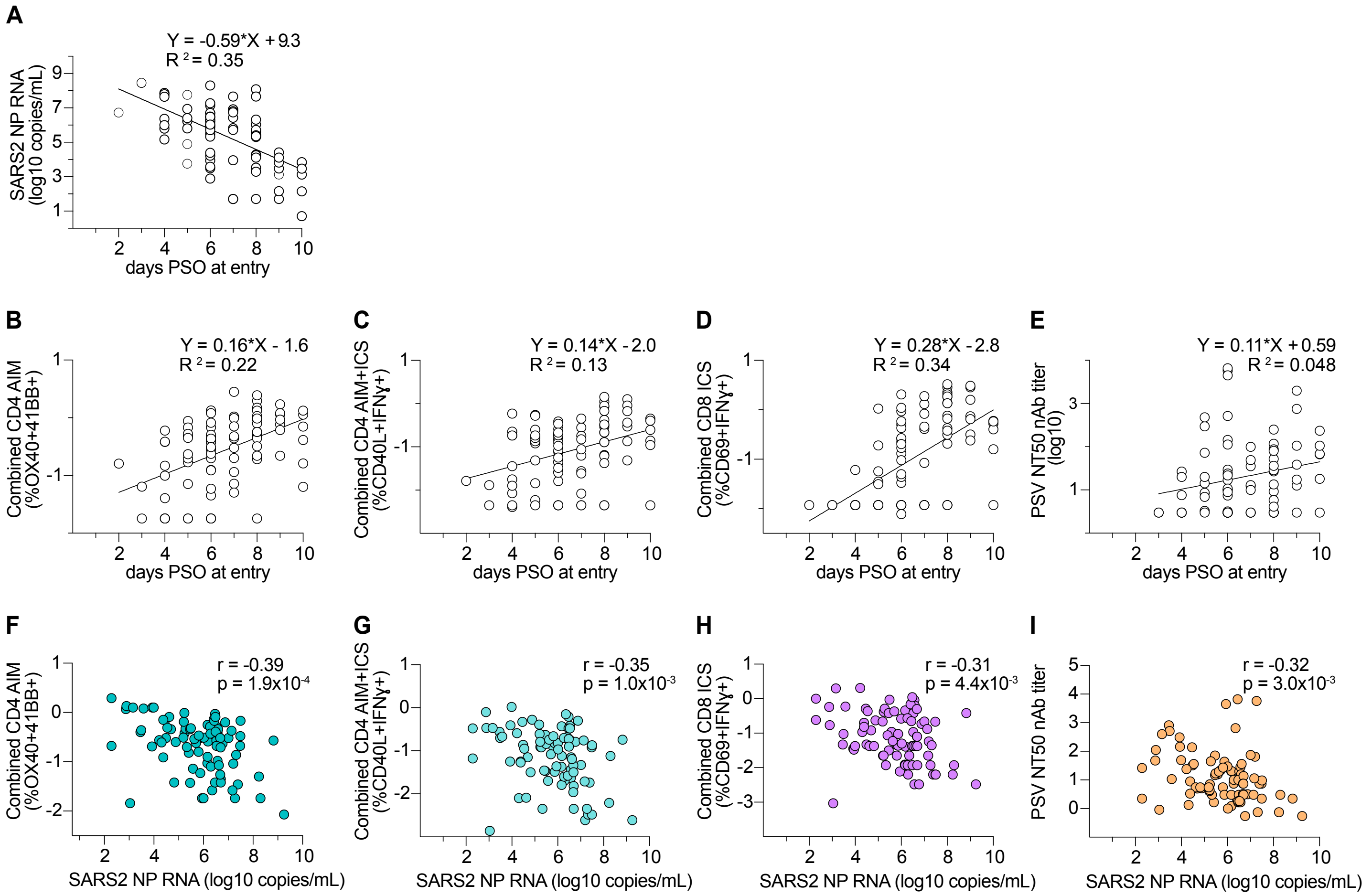


**Figure S7. Adjusted correlative relationships between virus-specific immune responses and upper airway SARS2 RNA levels.**
**A-E.** Log-linear regression models to account for variance in days from COVID-19 symptom onset to study entry (study day 0) for (A) SARS2 NP viral RNA, (B-C) SARS2-specific CD4 T cell responses, (D) SARS2-specific CD8 T cell responses, (E) and SARS2 nAb titers. Days post-symptom onset (PSO) at study day 0 values were plotted versus (A) SARS2 NP viral RNA or (B-E) immune responses at study day 0. Linear regression equations used for adjusted correlation analyses in **F-I** are shown. All immune response values in B-E were transformed to $\log_{10}$. **F-I.** Correlation analyses as in **Fig. 5A-D** for SARS2 NP RNA and SARS2-specific T cell responses by (**F**) CD4 $AIM^+$, (**G**) CD4 IFNγ, (**H**) CD8 IFNγ, and (**I**) nAb titers. In **F-I**, adaptive immune response values were adjusted to day 6 PSO using the related log-linear regression lines shown in **B-E** and plotted versus the corresponding adjusted day 6 PSO SARS2 NP RNA values generated from **A**. All correlations by two-tailed nonparametric testing; r = Spearman's rank correlation coefficient. **Related to Figure 5.**
